# SnoRNA guide activities – real and ambiguous

**DOI:** 10.1101/2021.07.15.452550

**Authors:** Svetlana Deryusheva, Gaëlle J.S. Talross, Joseph G. Gall

## Abstract

In eukaryotes, rRNAs and spliceosomal snRNAs are heavily modified posttranscriptionally. Pseudouridylation and 2’-*O*-methylation are the most abundant types of RNA modifications. They are mediated by modification guide RNAs, also known as small nucleolar (sno)RNAs and small Cajal body-specific (sca)RNAs. We used yeast and vertebrate cells to test guide activities predicted for a number of snoRNAs, based on their regions of complementarity with rRNAs. We showed that human SNORA24 is a genuine guide RNA for 18S-Ψ609, despite some non-canonical base-pairing with its target. At the same time, we found quite a few snoRNAs that have the ability to base-pair with rRNAs and can induce predicted modifications in artificial substrate RNAs, but do not modify the same target sequence within endogenous rRNA molecules. Furthermore, certain fragments of rRNAs can be modified by the endogenous yeast modification machinery when inserted into an artificial backbone RNA, even though the same sequences are not modified in endogenous yeast rRNAs. In *Xenopus* cells a guide RNA generated from scaRNA, but not from snoRNA, could induce an additional pseudouridylation of U2 snRNA at position 60; both guide RNAs were equally active on a U2 snRNA-specific substrate in yeast cells. Thus, posttranscriptional modification of functionally important RNAs, such as rRNAs and snRNAs, is highly regulated and more complex than simply strong base-pairing between a guide RNA and substrate RNA. We discuss possible regulatory roles for these unexpected modifications.

## INTRODUCTION

Small nucleolar RNAs (snoRNAs) are abundant non-coding nuclear RNAs. The vast majority of these RNAs function as guide RNAs to mediate site-specific post-transcriptional 2’-*O*-ribose methylation and pseudouridylation of other RNA molecules. Their specificity is determined by the so-called antisense elements (ASEs) – short sequences complementary to substrate RNAs. In the case of 2’-*O*-methylation, ASEs are located upstream of the D/D’ box (CUGA) motif in box C/D snoRNAs; exactly the fifth nucleotide from the D-box sequence determines the position of 2’-*O*-methylation in a substrate RNA. Pseudouridylation is mediated by box H/ACA snoRNAs. The ASEs in this type of modification guide RNA are formed by internal loops in 5’ and 3’ stems associated with box H (ANANNA) and box ACA, respectively; these loops are usually called pseudouridylation pockets. Typically, the distance from the box H or box ACA to the position of the target uridine inside the pocket is 15±1 nucleotide (reviewed in Bachellerie et al. 2002; Yu et al. 2005; Watkins and Bohnsack 2012). These are the basic rules for functional modification guide RNAs, yet they are neither sufficient nor absolutely essential (Deryusheva and Gall 2018).

Pseudouridylation and 2’-*O*-methylation are the most abundant and functionally important post-transcriptional modifications in eukaryotic rRNAs and spliceosomal snRNAs (Yu et al. 1998; Dönmez et al. 2004; Lapeyre 2005; Liang et al. 2007, 2009; Jack et al. 2011). With a few exceptions, positioning of these modifications depends on the guide RNA activities. Recent studies identified transcriptome-wide spread of pseudouridines and 2’-*O*-methylated residues (Carlile et al. 2014; Schwartz et al. 2014; Dai et al. 2017). The patterns and levels of RNA modifications are highly regulated, and they may be significantly modulated under different physiological and pathological conditions (Wu et al. 2011; Schwartz et al. 2014; Krogh et al. 2016; Sharma et al. 2017; Taoka et al. 2018).

Usually, computational or visual search for regions of complementarity between guide and substrate RNAs is used to predict specific targets for modification guide RNAs. However, such predictions may turn out to be false when they are tested experimentally (Xiao et al. 2009; Deryusheva and Gall 2018; Deryusheva et al. 2020). In a number of studies of eukaryotic snoRNA modification guide activities, *in vitro* reconstitution systems showed robust efficiency using cell nuclear extracts (Jády and Kiss 2001; Ma et al. 2005; Deryusheva and Gall 2009; Xiao et al. 2009; Deryusheva and Gall 2013) or recombinantly expressed and purified protein complexes (Kelly et al. 2019; Yang et al. 2020). At the same time, some snoRNAs that are inactive in cell free *in vitro* assays can function efficiently as modification guide RNAs in living cells (Deryusheva and Gall 2013, 2019b).

In our recent analysis of human and *Xenopus* snoRNA sets we found snoRNAs with regions complementary to 18S and 28S rRNAs at positions that have never been found modified in any species (Deryusheva et al. 2020). These postulated ASEs for unmodified positions were both evolutionarily conserved and species- or taxon-specific. We focus here on experimental verification of the predicted modification activities at unexpected positions. To our surprise a subset of tested snoRNAs could mediate predicted modifications in artificial substrate RNAs but did not do so in endogenous rRNAs. We hypothesize that some domains in rRNAs are not accessible for post-transcriptional modifications or become modified only under certain conditions such as stress. Alternatively, these domains may trigger RNA degradation when they are exposed for modification; that is, the modified RNA molecules are undetectable because they are unstable.

## RESULTS

### Human SNORA24 is a genuine guide RNA for position 609 in 18S rRNA

Since its discovery, human SNORA24 has been assigned to the known pseudouridine at position 609 in 18S rRNA (Kiss et al. 2004). Yet, in a later study SNORA24 activity on this position was not confirmed using yeast nuclear extract (Xiao et al. 2009). Since false-negative results are possible in this type of *in vitro* assay (Deryusheva and Gall 2013, 2019b) we repeated the test using living cells. First, we made a construct to express human SNORA24 *in vivo* in *S. cerevisiae*. For some reason this snoRNA did not accumulate in yeast. Therefore, we switched to a vertebrate cell system. In our previous study we performed comparative analysis of mammalian and amphibian snoRNAs and modification patterns of their predicted targets (Deryusheva et al. 2020). We found that the ASE for pseudouridilation of position 609 in 18S rRNA is missing from *Xenopus* SNORA24 and thus the equivalent position is not modified in *Xenopus*. This observation indicated that human SNORA24 might have modification activity on 18S rRNA at position 609. To verify this guide function of human SNORA24, we took advantage of the differences between human and *Xenopus* 18S rRNA modifications and transfected *X. laevis* cell line XTC with a human SNORA24 expression construct. Predictably, human SNORA24, when expressed in XTC cells, induced pseudouridylation of *Xenopus* 18S rRNA at the position equivalent to 609 in human 18S rRNA (Figure 1A). Thus, these results provide strong evidence that human SNORA24 is a genuine guide RNA for pseudouridylation of position 609 in 18S rRNA. Furthermore, we demonstrated again that *in vitro* reconstitution assays are less compatible with modification guide RNA activities than methods based on living cells.

**Figure 1.**
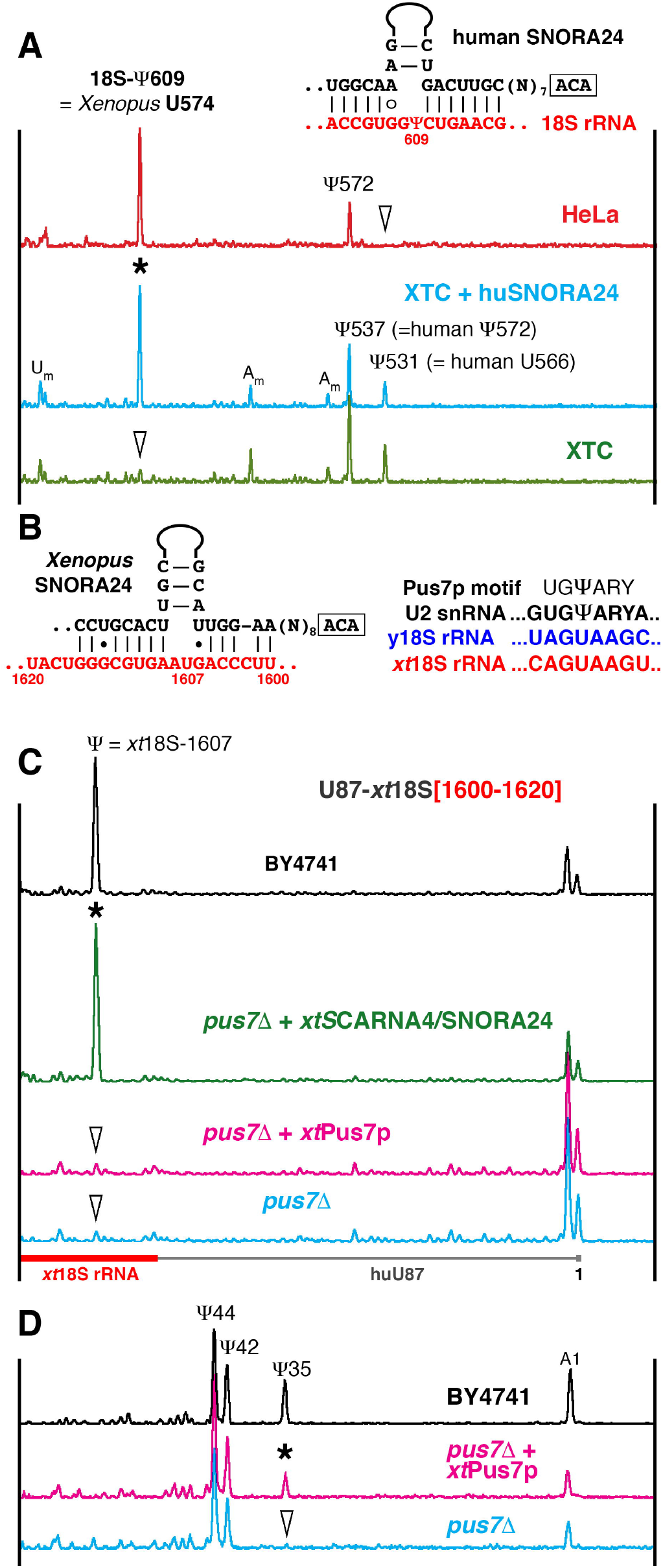
Testing SNORA24 modification guide activities. (A) Mapping of pseudouridines in 18S rRNA from human HeLa cells (red trace) and *Xenopus laevis* XTC cells (green and blue traces) using fluorescent primer extension. Peaks corresponding to reverse transcription termination at modified positions are indicated. Arrowheads point to unmodified positions. Ectopic expression of human SNORA24 in XTC cells induces pseudouridylation of *Xenopus* 18S rRNA at position 574, equivalent to human 18S-609 (blue trace, star). Postulated base-pairing between human SNORA24 and 18S rRNA shown in the top right corner. (B) Predicted base-pairing between *Xenopus* SNORA24 and 18S rRNA (left) and Pus7p recognition motif (right). (C) Modification of the artificial substrate RNA specific for *xt*SNORA24 (U87-*xt*18S[1600-1620]) in wild type BY4741and *pus7Δ* mutant yeast strains. The position corresponding to *xt*18S-1607 in the artificial substrate RNA is pseudouridylated in wild type (black trace) and in the *pus7Δ* strain that expresses *xt*SNORA24 (green trace, star). This position is not modified in the *pus7Δ* strain itself (blue trace, arrowhead) and in the *pus7Δ* strain that expresses *Xenopus* Pus7p (pink trace, arrowhead). Note, that in a fragment of 18S rRNA [1600-1620], yeast Pus7p recognizes a sequence different from the consensus UGΨAR (Urban et al. 2009; Carlile et al. 2014; Schwartz et al. 2014): it contains A instead of unvarying U at -2 position, AGΨAA (B). (D) Pseudouridylation of yeast U2 snRNA in wild type (black trace) and mutant *pus7Δ* strains (blue and pink traces). Expression of *Xenopus* Pus7p in the *pus7Δ* yeast strain rescues pseudouridylation of U2 snRNA at position 35 (pink trace, star).

### Artificial substrate RNAs, but not endogenous rRNAs undergo modification

*Xenopus* SNORA24, which lacks the pseudouridylation pocket for 18S-574 (equivalent to human 18S-609), shows complementarity to 18S rRNA at position 1607 (equivalent to human 18S-1649) (Figure 1B). Yet, neither position 574 nor 1607 is modified in *Xenopus* 18S rRNA (Deryusheva et al. 2020). To test *Xenopus* SNORA24 activity, we made a construct to express this guide RNA in yeast cells. As with human SNORA24, *Xenopus* SNORA24 did not express in yeast. Therefore, we designed a chimeric guide RNA construct from *Xenopus* SCARNA4 in which the 5’ terminal domain with a pseudouridylation pocket for position 41 of U2 snRNA was replaced with the 3’ terminal domain of *Xenopus* SNORA24. This SNORA24-SCARNA4 chimeric RNA was expressed efficiently in yeast. Because the yeast 18S rRNA sequence at the predicted *Xenopus* SNORA24 target position is slightly different from that of *Xenopus* 18S rRNA, an additional construct was made to assay *Xenopus* SNORA24 modification activity in yeast. This construct was used to express a fragment of vertebrate 18S rRNA as an artificial substrate RNA. Intriguingly, when this construct was expressed in the wild type yeast strain, the SNORA24 substrate RNA became modified at the tested position even without co-expression of exogenous guide RNA (Figure 1C, top black trace).

We screened several yeast mutant strains deficient for different pseudouridine synthases and found that yeast Pus7p was responsible for this modification (Figure 1C, blue trace), even though the modified sequence does not contain the canonical motif recognized by yeast Pus7p (Figure 1B). It is important to clarify here that yeast (Figure 1C, black trace), but not *Xenopus*, Pus7p (Figure 1C, pink trace) can modify that sequence. It seems that Pus7p synthases from different species, like Pus1p itself (Behm-Ansmant et al. 2006), differ in their activities on certain RNA substrates. In these experiments, modification of yeast U2 snRNA at position 35 served as an internal control for Pus7p modification activity (Figure 1D). Ultimately, in the mutant *pus7Δ* strain the chimeric SNORA24-SCARNA4 guide RNA could induce pseudouridylation of the artificial substrate RNA at the position corresponding to *Xenopus* 18S-1607 (Figure 1C, green trace). Thus, we observed that both *Xenopus* SNORA24 and yeast Pus7p can recognize and modify their target sequence when inserted in an artificial substrate RNA. However, they do not do so in endogenous rRNA molecules: neither position 1607 in *Xenopus* 18S rRNA (*xt*SNORA24 substrate) nor the equivalent position in yeast 18S rRNA (yeast Pus7p substrate) is modified.

The modification of the SNORA24 substrate RNA by yeast Pus7p is not the only case in which the endogenous modification machinery is active on an artificial substrate RNA but not on endogenous yeast rRNA. Another example is a potential substrate for one of the two copies of *X. tropicalis* SNORA15 (Supplemental Figure S1). Here again, no modification is found at the predicted target position in 18S rRNA from any species. Although the 18S rRNA sequence surrounding this position is extremely conserved, overexpression of *xt*SNORA15 in yeast cells did not induce modification of yeast 18S rRNA at position 1136. At the same time, when we expressed the corresponding artificial substrate RNA alone as a negative control, it became modified. That is, the endogenous yeast modification machinery recognizes and modifies this substrate RNA, but not the same sequence within endogenous yeast 18S rRNA. Unfortunately, in this case we could not identify the corresponding yeast guide RNA or stand-alone pseudouridine synthase, nor could we further test *xt*SNORA15 activity.

In addition to *xt*SNORA24 and yeast Pus7p, we identified several other vertebrate snoRNAs that were functional *in vivo* on artificial substrate RNAs, but not on endogenous rRNAs. One such snoRNA is the orphan vertebrate SNORA18. It has an evolutionarily conserved ASE that can base-pair with 18S rRNA to convert U1720 to pseudouridine (Figure 2A). Yet, this position is not modified in any vertebrate species tested so far (Deryusheva et al. 2020). When we expressed SNORA18 along with the corresponding artificial substrate RNA in yeast cells, the artificial substrate RNA became pseudouridylated at the predicted position (Figure 2B, pink trace). However, like vertebrate 18S rRNA, yeast 18S rRNA was not modified by SNORA18, even if its ASE was yeast-optimized and it was then overexpressed (Figures 2A, 3A, C-to-U mutant, blue traces).

**Figure 2.**
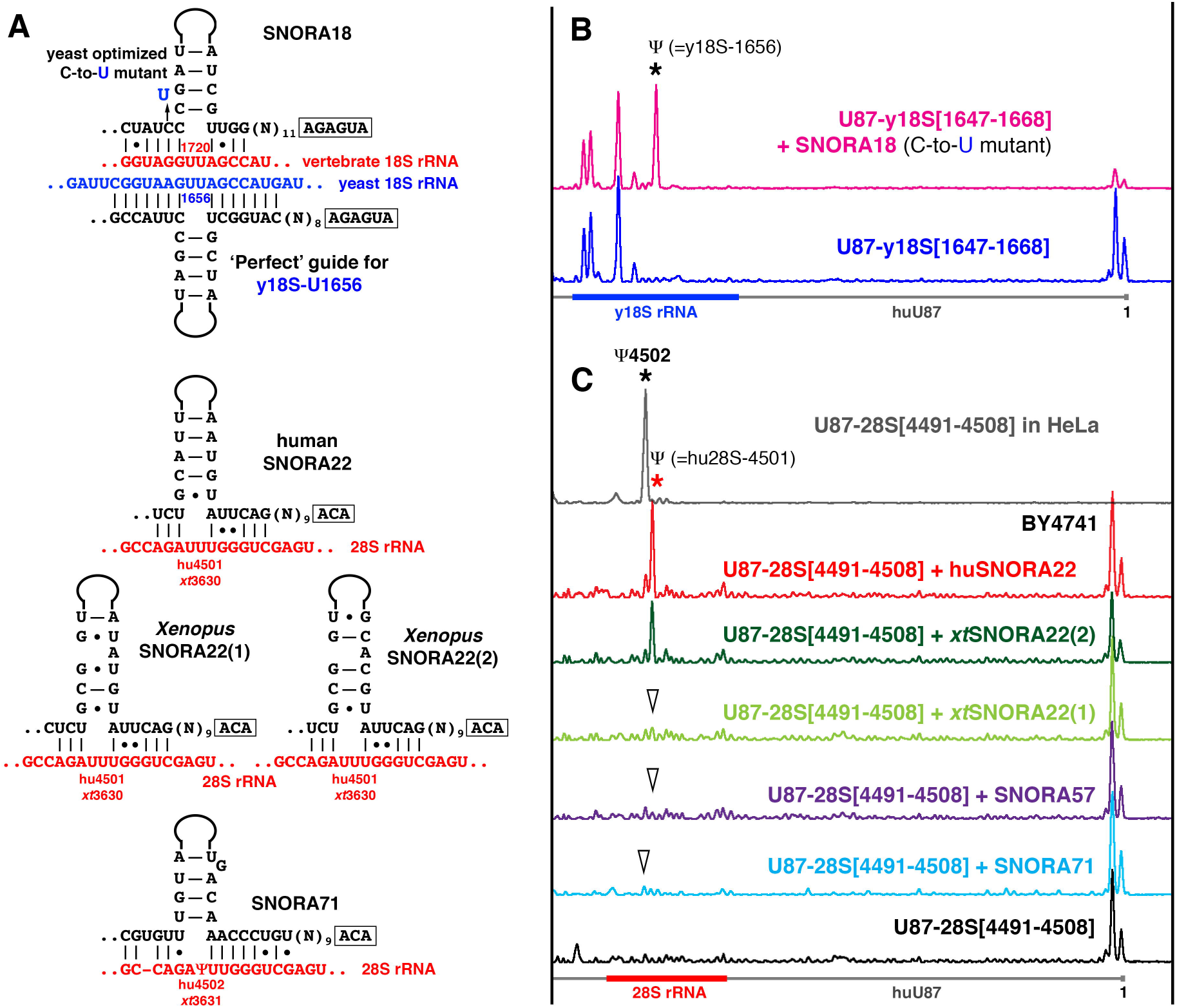
(A) Predicted base-pairing of SNORA18, SNORA22, and SNORA71 with 18S rRNAs at position 1720 and with 28S rRNA at positions 4501 and 4502, respectively. (B, C) Pseudouridylation of artificial substrate RNAs in yeast cells. (B) Yeast optimized SNORA18 (C-to-U-mutant) can induce predicted pseudouridylation in a fragment of yeast 18S rRNA [1647-1668 nt] inserted in U87 RNA (pink trace, star). Expression of the artificial substrate RNA U87-y18S[1647-1668] alone used as a negative control (blue trace). (C) When expressed in yeast cells, human SNORA22 (red trace, red star) and *Xenopus* SNORA22(2) (dark green trace) can induce pseudouridylation of position 4501 in a 28S rRNA fragment [4491-4508 nt] inserted in U87 RNA. *Xenopus* SNORA22(1) (light green trace, arrowhead) and SNORA57 (dark blue trace, arrowhead) do not show modification activity on the same substrate RNA. SNORA57 activity on 28S-4501 was predicted by Kehr et al. (Kehr et al. 2014); for the proposed base-pairing see Supplemental Figure 1. Note that two copies of *Xenopus* SNORA22 – functional and non-functional – differ only in their upper stem structures. In human, 28S rRNA position 4502 is pseudouridylated, and this position becomes modified in the artificial substrate RNA expressed in HeLa cells (top grey trace, black star). SNORA71 can base-pair with 28S rRNA at position 4502 (A). However, when it was tested in yeast, SNORA71 could not induce this modification in the artificial substrate RNA (light blue trace, arrowhead).

Also, we predicted that position 4501 in 28S rRNA should be a target for vertebrate SNORA22 (Figure 2A). In the original study of human 28S rRNA, pseudouridine was erroneously assigned to this position (Ofengand and Bakin 1997), whereas, in fact, position 4502 is pseudouridylated (Carlile et al. 2014; Taoka et al. 2018; Deryusheva et al. 2020). No guide RNA has been assigned so far for position 4502. When we tested SNORA22 modification activity in yeast cells, human SNORA22 and one of two copies of *Xenopus* SNORA22 could modify position 4501 in a corresponding artificial substrate (Figure 2C, red and dark green traces). However, when the same artificial substrate was expressed in human HeLa and *Xenopus* XTC cell lines, only Ψ4502 was detected (Figure 2C, top grey trace). Similar examples were already reported (Deryusheva and Gall 2013; Deryusheva et al. 2020). Nevertheless, we should emphasize here that in certain cases snoRNAs with quite strong postulated base-pairing between their ASEs and target RNAs could not modify endogenous RNAs nor could they modify artificial substrate RNAs (Figure 2C, Supplemental Figure S1; Deryusheva and Gall 2017, 2019a; Deryusheva et al. 2020).

### Heat shock does not induce 18S rRNA pseudouridylation

All snoRNAs that functioned on artificial substrates but not on endogenous rRNAs have somewhat imperfect base-pairing between their ASEs and the predicted target sequences (Figures 1B, 2A). For example, yeast Pus7p recognized and modified a sequence that is different from the canonical Pus7p-specific motif (Figure 1B). We hypothesized that such unusual activities might play regulatory roles and corresponding modifications might be induced on endogenous RNAs by stress. In fact, under stress conditions yeast snR81 interacts with a slightly diverged sequence to mediate stress-inducible modification of U2 snRNA (Wu et al. 2011). It has been shown that in response to heat shock, Pus7p induces RNA pseudouridylation at certain positions in U2 snRNA and mRNAs (Wu et al. 2011; Schwartz et al. 2014). However, pseudouridylation of 18S rRNA at position 1585, a modification we expected to be catalyzed by yeast Pus7p, was not reported for yeast exposed to a short one-hour heat shock at 45°C (Schwartz et al. 2014; Begik et al. 2021). We wondered if newly synthesized rRNA will gain additional modifications upon long exposure to heat. We compared pseudouridylation patterns of the 3’-terminal 400 nucleotides in yeast 18S rRNA isolated from wild type cells cultured at 30°C and at 37°C for over 24 hrs. We did not detect any differences between these samples (Figure 3A). That is, neither heat shock nor growth at high temperature induced yeast Pus7p activity on 18S-1585.

**Figure 3.**
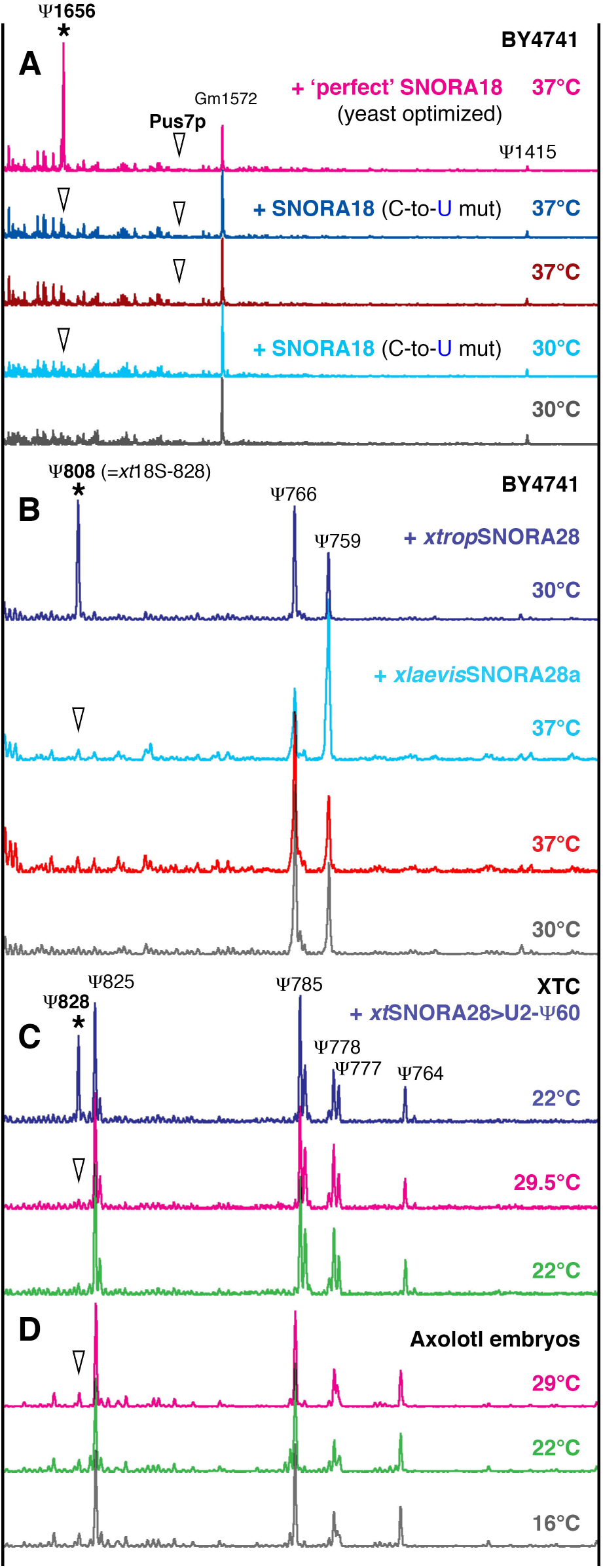
18S rRNA modification under normal and high temperature conditions in yeast (A, B) and amphibian cells (C, D). (A) Yeast optimized SNORA18 (C-to-U mutant, depicted in Figure 2A) does not induce pseudouridylation of yeast 18S rRNA at position 1656 (equivalent to human 18S-1720) in the BY4741 strain grown at 30°C, nor can it do so at 37°C (blue traces). ‘Perfect’ SNORA18, a ‘perfect guide’ RNA for position 1656 in yeast 18S rRNA made from SNORA18 (depicted in Figure 2A), induces pseudouridylation of the target position both at 30°C (not shown) and 37°C (top pink trace, star). Note there is no Pus7p activity on position 1585 in 18S rRNA both at 30°C and 37°C. Positions that were expected to be modified are indicated with arrowheads. (B) Yeast cells grown at 37°C do not induce pseudouridylation activity of *X. laevis* SNORA28 on yeast 18S rRNA at position 808 (equivalent to *Xenopus* 18S-828). Note the increased modification level at position 759, the other target of SNORA28 (light blue trace), as compared to controls (grey and red traces). *X. tropicalis* SNORA28 is functional on position 808 (top dark blue trace, star). (C, D) *X. laevis* XTC cells (C) and axolotl embryos (D) grown at elevated temperatures do not have altered pseudouridylation patterns of 18S rRNA. (C) When snoRNA with a functional pseudouridylation pocket for 18S-828 (*xt*SNORA28>U2-Ψ60, blue trace) is expressed in XTC cells, this position is modified (star). Unmodified position 828 is indicated with arrowheads.

In the same 400-nt 3’-terminal region of 18S rRNA, we predicted the potential target for SNORA18. We grew yeast cells transformed with the expression construct for a yeast optimized variant of SNORA18 (Figure 2A, C-to-U mutant) at 37°C to a high cell density (OD_600_ ∼ 5-6). These growing conditions did not induce modification activity of the yeast optimized SNORA18 on yeast 18S rRNA at position 1656 (Figure 3A, dark blue trace). To demonstrate that this position in helix 44 of yeast 18S rRNA could be pseudouridylated, we extended the SNORA18 ASE and changed two non-canonical U-G base pairs to C-G (Figure 2A, ‘perfect’ guide). This perfect guide RNA was functional on yeast 18S rRNA (Figure 3A, top pink trace), indicating that this position is modifiable.

Finally, we tested whether elevated temperatures could induce activity of *X. laevis* SNORA28 on 18S rRNA at position 828 (equivalent to position 866 in human and position 808 in yeast 18S rRNAs). *X. laevis* SNORA28 contains the corresponding ASE, yet we observed the lack of this modification in *X. laevis* 18S rRNA. In fact, this ASE is somewhat imperfect in *xl*SNORA28. When this snoRNA was tested in the yeast cell system, it showed modification activity on a fragment of rRNA inserted in an artificial backbone RNA, but could not modify yeast 18S rRNA at position 808 (Deryusheva et al. 2020). We noticed that the lack of pseudouridylation of 18S rRNA at position 828 in three amphibian species, *X. laevis*, axolotl and newt, but not in *X. tropicalis* (much warmer climate), correlated with the environmental conditions in which these species normally live (Deryusheva et al. 2020). We speculated that higher temperature might enhance modification activity of the imperfect pseudouridylation pocket of *xl*SNORA28.

We grew yeast cells transformed with *xl*SNORA28 expression constructs at 37°C. In these conditions *xl*SNORA28 still could not induce pseudouridylation of yeast 18S-U808 (Figure 3B). Then we cultured the *X. laevis* cell line XTC at 29.5°C; importantly, above this temperature these cells start dying in a few hours. 18S rRNA from the heat stressed XTC cells did not have additional pseudouridine at position 828, the potential target for *xl*SNORA28 (Figure 3C). Therefore, when *X. tropicalis* SNORA28 with a functional 18S-828 pseudouridylation pocket was expressed in XTC cells, *xl*18S rRNA became pseudouridylated at position 828 under normal conditions (Figure 3C, *xt*SNORA28>U2-Ψ60; Deryusheva and Gall 2019a; Deryusheva et al. 2020).

Additionally, we analyzed 18S rRNA from axolotl embryos raised at different temperatures between 16° and 29°C until they were about to hatch; embryos that developed at 29°C never hatched. As we mentioned above, in this species 18S rRNA is missing pseudouridylation at position 828. Axolotls are more temperature sensitive than frogs, but their 18S rRNA modification patterns still showed no pseudouridylation at position 828 in the heat shock conditions (Figure 3D). Thus, taken together our data suggest that heat does not facilitate pseudouridylation activity of imperfect snoRNAs on 18S rRNA in yeast and vertebrate cells.

### Assisted modification activity

SNORA57 was originally identified as a guide RNA for pseudouridylation of position 1004 in human 18S rRNA (Schattner et al. 2006). The configuration of the pseudouridylation pocket for this modification is not canonical: it has three target nucleotides within the pocket instead of two (Figure 4A). We tested SNORA57 modification activity on yeast 18S rRNA, which does not have pseudouridine at the equivalent position 947 (Figure 4B, black trace). When we expressed human SNORA57 in yeast cells, we did not detect pseudouridylation of position 947 in yeast 18S rRNA (Figure 4B, red trace). Perhaps the yeast 18S rRNA sequence is slightly diverged at the target position, which makes SNORA57 nonfunctional on endogenous rRNA. Therefore, we made a yeast optimized version of SNORA57 (Figure 4A, U-to-C mutation) and expressed it in yeast cells. The yeast optimized SNORA57 did induce pseudouridylation of position 947 in yeast 18S rRNA. Additionally, it induced pseudouridylation at nearby position 989 equivalent to human 1046 (Figure 4B, blue trace, stars).

**Figure 4.**
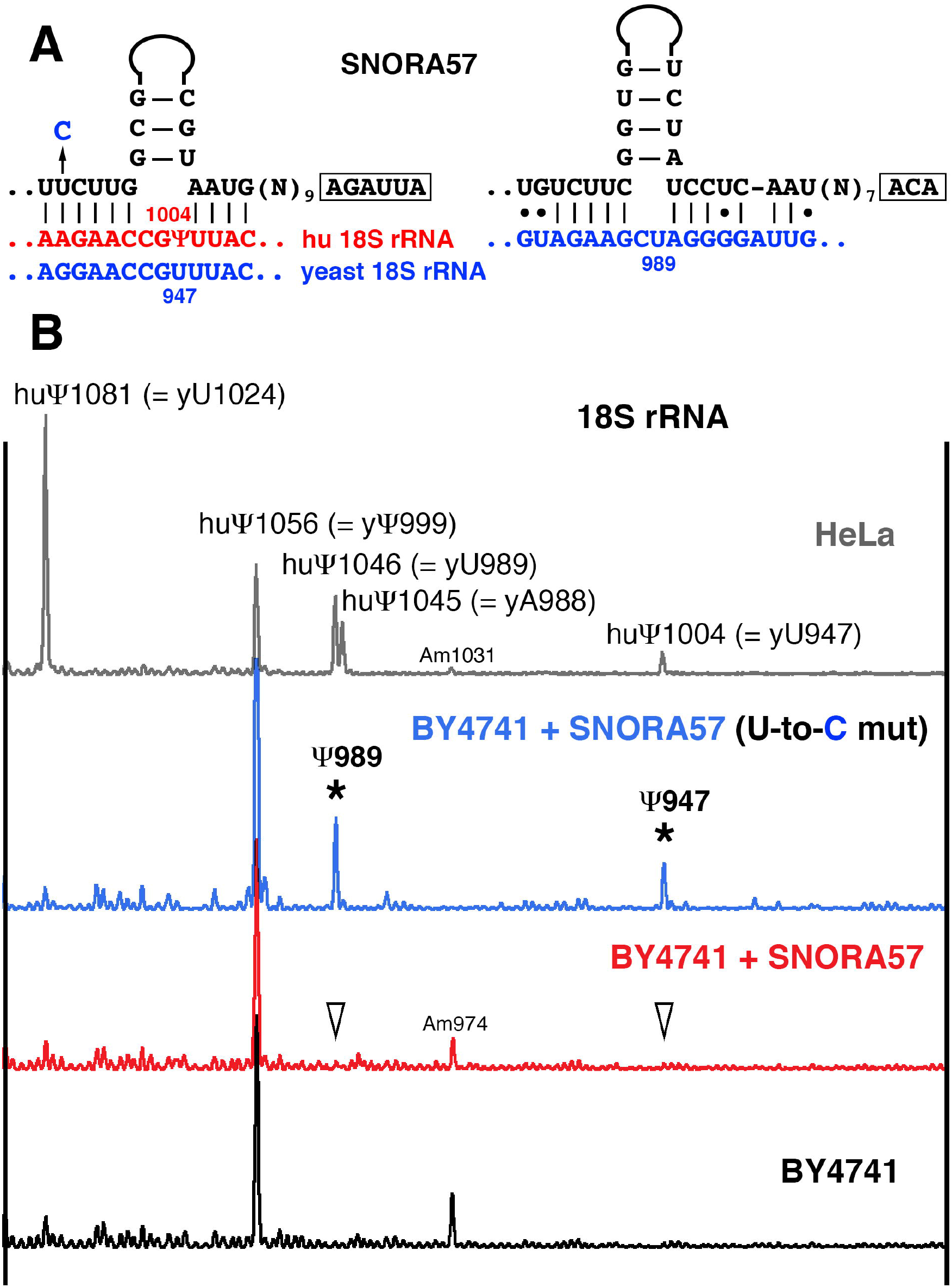
SNORA57 guide activities on yeast 18S rRNA. (A) Postulated base-pairing of SNORA57 with human (red) and yeast (blue) 18S rRNA. To make a yeast optimized version of SNORA57 an indicated U-to-C mutation was introduced in the ASE of the 5’ terminal pseudouridylation pocket. (B) Mapping pseudouridines in human 18S [968-1086 nt] (HeLa, top grey trace) and the equivalent region of yeast 18S rRNA (BY4741, black, red and blue traces). Normally, yeast 18S rRNA has only Ψ999 in this region (black trace). Expression of vertebrate SNORA57 did not change the modification pattern of yeast 18S rRNA (red trace, arrowheads indicate predicted target positions for SNORA57). Yeast optimized SNORA57 with the U-to-C mutation could induce pseudouridylation of yeast 18S rRNA at two positions, 947 and 989 (blue trace, stars).

Recently, we and others detected two novel pseudouridines 1045 and 1046 in 18S rRNA from humans and various vertebrate species (Carlile et al. 2014; Stanley et al. 2016; Taoka et al. 2018; Deryusheva et al. 2020). We showed that the 3’ terminal pseudouridylation pocket in SNORA57 can mediate pseudouridylation of these positions in vertebrate 18S rRNA (Deryusheva et al. 2020). The same ASE must have base-paired with yeast 18S rRNA and mediated pseudouridylation at position 989. However, the proposed base-pairing is imperfect (Figure 4A), and this weak pseudouridylation pocket is functional only in conjunction with much stronger interaction within the other pocket. In short, the modification activity of a weak pseudouridylation pocket can be enhanced by strong base-pairing elsewhere in the molecule.

### Another layer of complexity in higher eukaryotes

In yeast, a single guide RNA, snR81 modifies both 25S rRNA and spliceosomal U2 snRNA (Ma et al. 2005; Wu et al. 2011). Yet, in higher eukaryotes, a specialized class of modification guide RNAs is involved in endogenous spliceosomal U snRNA modification. These are the so-called scaRNAs, concentrated in the nuclear Cajal bodies (CBs). In our previous study we showed that both snoRNAs and scaRNAs can modify certain positions in rRNAs (Deryusheva and Gall 2019a). However, it is still an open question whether scaRNAs and snoRNAs are interchangeable in the case of endogenous snRNA modification in higher eukaryotes.

To assess the ability of a typical snoRNA to modify U2 snRNA in a vertebrate cell system we took advantage of the differences between *Xenopus* and mammalian U2 snRNA modification patterns. Pseudouridine at position 60 has been reported in human (Figure 5B, bottom grey trace) and rodent U2 snRNA (Deryusheva et al. 2012; Deryusheva and Gall 2017), although in *Xenopus* this position is not modified (Figure 5B, green trace). We generated artificial U2-Ψ60 guide RNAs from *X. tropicalis* SCARNA4 and SNORA28 by ASE replacement in their U2-Ψ41 and 18S-Ψ778 pockets, respectively. Importantly, the configuration of U2-Ψ60 pockets was identical in both constructs (Figure 5A). These two artificial guide RNAs, which could target U2-U60 for pseudouridylation, were expressed in the *Xenopus* XTC cell line. In these experiments, only guide RNA made from scaRNA (SCARNA4>U2-Ψ60) could induce pseudouridylation of *Xenopus* U2 snRNA at position 60 (Figure 5B). As a control these guide RNAs were expressed in yeast cells along with the corresponding artificial substrate RNA (U87-U2[51-68]); SNORA28>U2-Ψ60 was functional on the substrate RNA even if its ASE was shortened (Figure 5C). Since the second pseudouridylation pocket specific for *Xenopus* 18S-828 remained intact in SNORA28>U2-Ψ60, it served as an internal control for this guide RNA function in XTC cells. As expected, 18S rRNA became modified at position 828 when SNORA28>U2-Ψ60 was expressed in XTC cells (Figure 3C, top dark blue trace). Thus, our data demonstrate a much higher complexity of posttranscriptional modification on endogenous RNAs in living cells than one would anticipate based on *in vitro* RNA systems.

**Figure 5.**
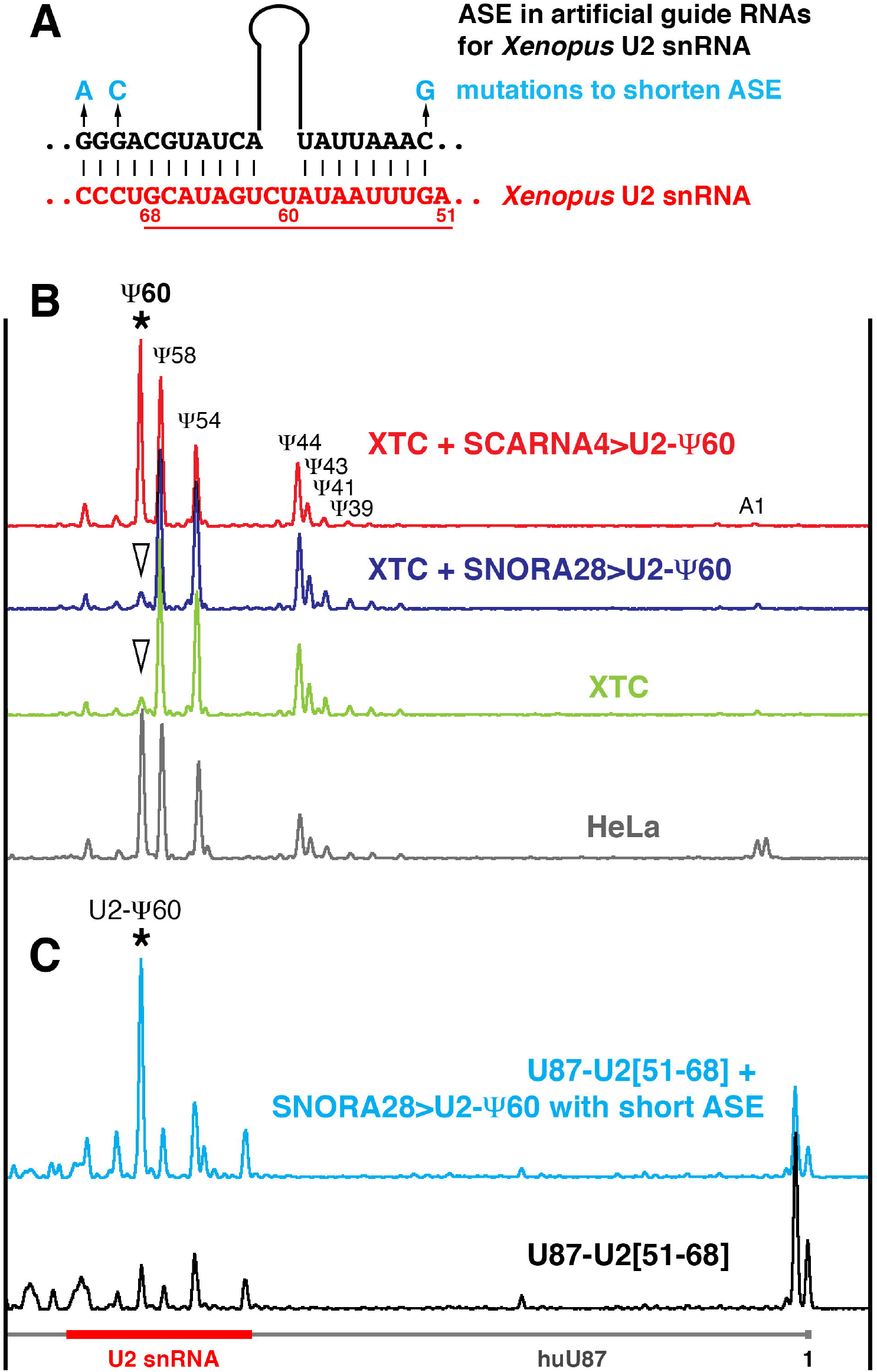
(A) A pseudouridylation pocket generated in *xt*SCARNA4 and SNORA28 to make artificial guide RNAs for *Xenopus* U2 snRNA modification at position 60 (SCARNA4>U2-Ψ60 and *xt*SNORA28>U2-Ψ60). Mutations that shorten the ASE are indicated in blue; a fragment of U2 snRNA inserted in a U87-based artificial substrate RNA is underlined. (B) Mapping pseudouridines in human and *Xenopus* U2 snRNA. Normally, human U2 snRNA (HeLa, grey trace) is pseudouridylated at position 60; *Xenopus* U2 snRNA is not modified at this position (XTC, green trace). Expression of SCARNA4>U2-Ψ60 (red trace, star) but not *xt*SNORA28>U2-Ψ60 (dark blue trace, arrowhead) induced U2 snRNA pseudouridylation at position 60 in the *Xenopus laevis* cell line XTC. (C) Pseudouridylation of the artificial substrate RNA containing a fragment of vertebrate U2 snRNA (U87-U2[51-68]) in yeast cells. Even with a short ASE, *xt*SNORA28>U2-Ψ60 is functional on U2 snRNA artificial substrate (blue trace, star).

## DISCUSSION

In this study we demonstrated that the process of posttranscriptional RNA modification in living cells is more complex than in *in vitro* systems. Using a *Xenopus* cell line (Figure 1A) instead of nuclear extracts (Xiao et al. 2009), we were able to verify modification guide RNA activity of human SNORA24 on 18S rRNA at position 609. At the same time, some predicted configurations of guide RNA-substrate RNA interactions remain nonfunctional even in living cells. For instance, in yeast cells, human SNORA19 did not modify yeast 18S rRNA at position 808 (equivalent to position 866 in human) (Xiao et al. 2009) nor did *Xenopus* SNORA19 substitute for missing SNORA28 activity on the equivalent position of 18S rRNA in *X. laevis* (Figure 3C; Deryusheva et al. 2020). Both results allow us to rule out SNORA19 modification activity on 18S-866.

Can we draw unequivocal conclusions by testing guide RNA activities in living cells? It’s complicated. Some snoRNAs that do not modify their predicted targets in endogenous rRNAs could function as modification guide RNAs on the same sequence inserted in an artificial RNA backbone (Deryusheva and Gall 2013; Deryusheva et al. 2020; this study). That is, the RNA content outside the targeted region itself can affect guide RNA activities. In fact, very subtle changes in the secondary structure of a substrate RNA can make it unmodifiable (Deryusheva and Gall 2019a). Those snoRNAs that show modification activity on artificial substrates but not on endogenous RNA targets usually contain somewhat imperfect ASEs. However, many fully functional snoRNAs have non-canonical base-pairing with their target RNAs (e.g. human SNORA24 tested in this study). Furthermore, some imperfection could even be evolutionarily conserved and essential for proper RNA modification pattern, especially in heavily modified regions where the positioning of modifications interfere with each other (Deryusheva and Gall 2018). Such interference can explain the absence of modification activities for some snoRNAs.

On the other hand, certain ASEs in snoRNAs might function as auxiliary ASEs to facilitate modification mediated by different ASEs. Extra base-pairing of ‘accessory guide’ with rRNA was found critical for some yeast snoRNAs function (van Nues and Watkins 2017). Similar additional base-pairing in vertebrate SNORD53, SNORD92 and *xt*SNORD88 might be needed for efficient modification (Supplemental Figure S1). Besides, these additional interactions might allow substrate RNA to slide easier into an alternative configuration of a functional modification guide RNA domain. That is what we presumably observed in the cases of vertebrate U92/SCARNA8 activities on U2 snRNA (Deryusheva and Gall 2017), and yeast optimized SNORA57 activity on yeast 18S rRNA (Figure 4).

Alternatively, non-functional RNA modification domains in snoRNAs could negatively regulate modification levels by competing with functional modification guide RNA ASEs for base-pairing within the same region. For instance, SNORD12 (non-functional domain) and SNORD48 (functional) complementary regions are overlapping. This might be the reason that 28S-Cm1868 is only fractionally modified by SNORD48 (Krogh et al. 2016; Taoka et al. 2018). Similarly, multiple overlapping interactions of SNORD53 and SNORD92 with 28S rRNA correlate with hypomethylated positions A3846 and C3848. In support of these alternative functions of ASEs in snoRNAs, unexpected physical interactions between box C/D snoRNAs and unmodified regions in rRNAs and snRNAs were captured by snoRNP CLIP analysis along with interactions of one modified rRNA site with multiple snoRNAs (Gumienny et al. 2017).

As we mentioned above, non-canonical ASEs are typical for snoRNAs that showed guide RNA modification activities on artificial substrates but not on the same sequences within endogenous RNA molecules. Such imperfect snoRNA interaction with substrate RNA is essential for stress inducible modification of U2 snRNA by yeast snR81 (Wu et al. 2011). It was previously noticed that pseudouridylation levels clearly correlate with environmental temperatures (Rajan et al. 2019; Deryusheva et al. 2020). However, we did not detect additional pseudouridines in 18S rRNA isolated from amphibian and yeast cells grown at elevated temperatures. Thus, exposure to heat could not induce unusual activities of SNORA18 and *xl*SNORA28 on 18S rRNA. On the other hand, some other types of stress might be involved in induction of RNA modifications mediated by imperfect yet potentially functional guide RNAs. In fact, in yeast U2 snRNA only one out of two inducible pseudouridines is formed in response to heat shock, and this isomerization is catalyzed by stand-alone pseudouridine synthases Pus7p; the other, snoRNA-dependent, inducible pseudouridine forms in response to nutrient stress (Wu et al. 2011). Another type of environmental stress, the filamentous growth, induces pseudouridylation of yeast U6 snRNA (Basak and Query 2014). Additional stress conditions that regulate RNA modification are still to be uncovered.

Another possibility is that these modifications occur only on mutated, misfolded or misassembled RNA. These abnormally modified RNA molecules might be targeted for degradation and therefore undetectable. Since we predicted and analyzed unusual pseudouridine synthase activities on 18S rRNA, we chose several yeast strains (*dom34Δ asc1Δ, dom34Δ hbs1Δ, ski7Δ hbs1Δ* and *ubc4*Δ generously provided by Melissa Moore) deficient for factors involved in 18S nonfunctional rRNA decay, or NRD (Limoncelli et al. 2017) for pilot experiments. We expressed SNORA18 in the listed mutant strains both under normal and heat shock conditions. In the mutant strains pseudouridylation of yeast 18S rRNA at positions 1656 (SNORA18 target), 1585 (yeast Pus7p) and 1136 (unknown yeast pseudouridine synthase, *xt*SNORA15) was not detected (data not shown). These negative results do not exclude the possibility that a novel 18S NRD-independent pathway of 18S rRNA quality surveillance might be triggered. This line of rRNA modification research is worth pursuing.

Thus, our data demonstrate that living cells are more proficient at supporting guide RNA-mediated RNA modifications than cell free *in vitro* systems. However, additional levels of regulation are observed *in vivo*. While ectopically expressed modification guide and substrate RNAs can interact in the cell to successfully position predicted modifications in the artificial substrates, modification of the same targets in endogenous rRNAs is tightly regulated. Abnormal modification of certain domains could be detrimental for cells. Indeed, induction of random pseudouridylation in *E. coli* 23S rRNA revealed critical regions whose hyper modification inhibits ribosome assembly (Leppik et al. 2017). In eukaryotes this regulation becomes incredibly complex. In yeast, specialized RNA helicases facilitate 2’-*O*-methylation of the particular positions in rRNAs (Aquino et al. 2021). In humans, Nopp140 chaperone interacts with both box C/D and box H/ACA snoRNPs and is required for concentration of both C/D and H/ACA scaRNAs in CBs. Yet Nopp140 is essential for 2’-*O*-methylation but not for pseudouridylation of spliceosomal snRNAs. The depletion of Nopp140 from HeLa cells causes dramatic reduction of 2’-*O*-methylation at all positions except for two in polymerase 2-transcribed snRNAs, but affects a very limited number of positions in rRNAs (Bizarro et al. 2021). Another RNA-binding protein, TDP-43, also contributes to 2’-*O*-methylation of U1 and U2 snRNAs. However, the positions that were not affected in the TDP-43KD differ from those in Nopp140KD cells (Izumikawa et al. 2019).

The function of CBs and scaRNAs in RNA modification is still not clear. In the fruit fly *Drosophila melanogaster*, the absence of CBs in coilin-null mutants has no effect on snRNA modification (Deryusheva and Gall 2009). *Drosophila* scaRNAs with proposed dual functions on snRNAs and rRNAs could recognize and modify rRNA fragments in yeast cell system (Deryusheva and Gall 2013), but not within endogenous rRNA in *Drosophila*. At the same time, typical scaRNAs when expressed ectopically in human or *Xenopus* cell lines can induce modification of vertebrate rRNAs at certain positions (Deryusheva and Gall 2019a). In *Xenopus* oocyte system *Drosophila* 2’-*O*-methylation guide RNA, scaRNA:MeU2-C28, both in a form of full-length scaRNA and mutated versions that lack CB-localization signals, can efficiently modify *in vitro* transcribed U2 snRNA (Deryusheva and Gall 2009, 2019b). Yet, efficient modification of endogenous U2 snRNA in vertebrate cells required canonical scaRNP to be involved both in pseudouridylation (Figure 5) and 2’-*O*-methylation (Deryusheva and Gall 2018; our unpublished observations). How can all these pieces of data fit a complete model for the regulated positioning of RNA modifications *in vivo*? Nowadays, the role of posttranscriptional modifications cannot be overstated. However, we are still in an early phase of understanding how RNA modification patterns of various RNA classes are established and regulated in different cell types, developmental stages, physiological and pathological conditions.

## MATERIALS AND METHODS

### Expression constructs for snoRNAs and artificial substrate RNAs

To express human SNORA24 and modified *X. tropicalis* SNORA28 and SCARNA4 in the *X. laevis* XTC cell line, fragments of these snoRNA host genes were amplified from genomic DNAs and cloned into the pCS2 vector. Overlap extension PCR was used to replace pseudouridylation pockets in SNORA28 and SCARNA4. Constructs for expression of vertebrate snoRNAs in yeast were generated by cloning corresponding snoRNA coding sequences into the YEplac181 vector, which contains a *GPD* promoter, an RNT1 cleavage site and an snR13 terminator (Huang et al. 2011). This vector was generously provided by Yi-Tao Yu, University of Rochester Medical Center. PCR-based mutagenesis was used to make yeast-optimized versions of vertebrate snoRNAs.

Artificial substrate RNA constructs were made as previously described (Deryusheva and Gall 2013, 2019a). In brief, human U87 scaRNA without the H/ACA domain was used as a backbone for insertions of various fragments of vertebrate and yeast rRNAs. Chimeric sequences were used to substitute for snR18 in the *EFB1* gene fragment cloned into the yeast p426Gal1 vector. For expression in vertebrate cells the insertions were subcloned into the pCS2 vector under the CMV promotor.

To express test RNAs ectopically, the listed constructs were introduced into yeast and *Xenopus* cells. Yeast “wild type” BY4741 or various mutant strains were transformed according to a standard lithium acetate protocol. *X. laevis* XTC cells were transfected using FuGene HD transfection reagent (Promega). Selective synthetic media containing either glucose or galactose as a sugar source were used to grow yeast cells. Heat shock conditions were 37°C for yeast and 29°C for amphibians.

The annotated protein coding sequence for *X. tropicalis* pseudouridine synthase Pus7 was amplified from total RNA using the OneTaq One-step RT-PCR kit (New England Biolabs). XmaI and XhoI restriction sites were added with oligonucleotides and the resulting fragment was cloned into the p415Gal1 vector.

### RNA extraction and ectopic expression analysis

RNA was extracted from control and transfected cells using the Trizol reagent in the case of vertebrate cells or hot acid phenol in the case of yeast cells. RNA was purified using the Direct-zol RNA miniprep kit (Zymo Research). Proper processing and expression levels of exogenous RNAs were analyzed using northern blotting as described (Deryusheva and Gall 2013).

### RNA modification analysis

To detect pseudouridines and 2’-*O*-methylated residues, we used fluorescent primer extension-based techniques as described (Deryusheva and Gall 2009; Deryusheva et al. 2012). RNA treated with CMC (N-cyclohexyl-N9- [2-morpholinoethyl] carbodiimide metho-p-toluene sulfonate) was used in reverse transcription reactions for pseudouridine mapping. A low concentration of dNTP was used to determine 2’-*O*-methylated positions. 6-FAM-labeled oligonucleotides specific for yeast and vertebrate U2 snRNA, 18S rRNA, and U87-based artificial substrate RNAs were described earlier (Deryusheva et al. 2012; Deryusheva and Gall 2013; Deryusheva et al. 2020). 6-FAM-labeled RT reaction products mixed with the Liz-500 Size Standard were separated on capillary columns using an ABI sequencing apparatus and a standard protocol for fragment analysis. GeneMapper 5 software (Applied Biosystems) was used to visualize and analyze the data.

## Acknowledgements

We are grateful to Yi-Tao Yu, Melissa J. Moore and Beverly Wendland for sharing yeast strains and vectors. This research was supported by the National Institute of General Medical Sciences of the National Institutes of Health (grant number R01 GM33397 to J.G.G.). J.G.G. is American Cancer Society Professor of Developmental Genetics.

## FIGURE LEGENDS

**Supplemental Figure S1**.

Potential base-pairing of snoRNAs with rRNAs at unmodified positions. rRNA fragments used in the artificial substrate RNAs are shown in red. For *Xenopus*-specific ASEs for unmodified positions, equivalent positions in human rRNAs are indicated in parentheses.

